# High-performance *Saccharomyces cerevisiae*-based biosensor for heavy metal detection

**DOI:** 10.1101/2021.12.13.472527

**Authors:** Cong Fan, Danli Zhang, Qiwen Mo, Jifeng Yuan

**Author notes:** These authors contributed equally to the experimental work (Cong Fan, Danli Zhang).

## Abstract

Heavy metals, i.e., Cu(II), are harmful to the environment. There is an increasing demand to develop inexpensive detection methods for heavy metals. Here, we developed a yeast biosensor with reduced-noise and improved signal output for potential on-site copper ion detection. The copper-sensing circuit was achieved by employing a secondary genetic layer to control the galactose-inducible (GAL) system in *Saccharomyces cerevisiae*. The reciprocal control of the Gal4 activator and Gal80 repressor under copper-responsive promoters resulted in a low-noise and ultrasensitive yeast biosensor for the copper ion detection. Furthermore, we developed a betaxanthin-based colorimetric assay, as well as 2-phenylethanol and styrene-based olfactory outputs for the copper ion detection. Notably, our engineered yeast sensor confers a narrow range switch-like behavior, which can give a “yes/no” response when coupled with betaxanthin-based visual phenotype. Taken together, we envision that the design principle established here might be applicable for developing other sensing systems for various chemical detections.

**IMPORTANCE:** The accumulation of heavy metals in the environment led to an imbalance with environmental and ecological impacts, and there is an increasing demand for inexpensive methods that permit on-site detection of heavy metals. In this study, a yeast biosensor with reduced-noise and improved signal output was developed. We used a betaxanthin-based colorimetric assay for monitoring the copper contamination in water samples, which confers a switch-like behavior with a “yes/no” response. In addition, we also explored the 2-phenylethanol and styrene-based outputs for copper detection, which might be useful for olfactory detection of heavy metals.

## INTRODUCTION

Heavy metals are natural-occurring chemicals that are present in the environment. However, due to massive industrialization, rapid urbanization and intensive agriculture, the accumulation of heavy metals in the environment led to an imbalance with environmental and ecological impacts (1). Some of the heavy metals (Cu, Mn, Zn and Co) are essential trace elements, playing an important role in maintaining the enzyme activities. For instance, copper is an essential trace element that is required for various cellular enzymes including cytochrome c oxidase, superoxide dismutase, and dopamine monooxygenase (2). Other heavy metals (Cd, Ag, and Hg) are not essential but interfere with the cellular metabolism, and disrupt biological functions. Nevertheless, when heavy metals are in excess amounts, they are all deleterious to living organisms (3, 4).

In the past years, techniques such as anodic stripping voltammetry (ASV) (5), graphite furnace atomic absorption spectrometry (6), and X-ray fluorescence spectrometry (7), have been developed for heavy metal analysis. However, all the above detection methods rely on the use of sophisticated instruments and highly qualified staff. There is a pressing need to develop alternative and inexpensive methods that permit on-site detection of metal ions. More recently, a microfluidic paper-based analytical device (mPAD) has been designed for quantifying metals in water, and the detection limit for copper and zinc can be as low as 0.1 ppm (8, 9). With the advancement of synthetic biology, there is an emerging interest in developing field-deployable “biosensors” for water quality monitoring (10). For instance, a cardiac cell-based biosensor has been employed for heavy metal detection (11). After exposure of cardiomyocytes to different heavy metal ions, there would be characteristic changes of beating frequency, amplitude and duration, which can be monitored by the light-addressable potentiometric sensor. In addition, a heavy metal monitoring bacterial system was also reported in *Escherichia coli* (12). The two-component system comprises membrane-associated sensor kinases such as ZraSR and CusSR with their cognate regulators in regulating the expression of ZraP and CusC to sense Zn(II) and Cu(II), respectively. However, the bacterial sensor had a very high background noise, and the signal output was only amplified several-fold upon the heavy metal exposure (12).

The first *Saccharomyces cerevisiae* biosensor that utilizes the copper-responsive promoter P_CUP1_ to drive the lacZ reporter for detecting Cu(II) by an amperometric method was reported two decades ago (13), however, the sensor could only measure the Cu(II) concentrations ranging from 0.5 to 2 mM. Since the first *S. cerevisiae* biosensor to detect Cu(II) was reported, a number of yeast biosensors for detecting heavy metals have been developed (14). For instance, a fluorescence-based sensing system for Cu(II) was designed via the P_CUP1_ controlled green fluorescent protein (GFP) expression (15) or luciferase (16). More recently, P_CUP1_ controlled expression of ADE5 and ADE7 in Δ*ade2* yeast strain to give the red pigment was also reported (17). However, all the above-mentioned systems rely on the leaky CUP1 promoter, which gives relatively high background noises (18, 19). In this study, synthetic biology design principles were applied to address the background noise commonly encountered by the CUP1-mediated sensing systems in yeast. We have reprogrammed *S. cerevisiae* into a low-noise, ultrasensitive and inexpensive device for Cu(II) detection without an additional requirement of equipment.

## MATERIALS AND METHODS

### Plasmid construction and yeast genome engineering

*E. coli* strain TOP10 was used for routine plasmid constructions. *S. cerevisiae* strain BY4741 was obtained from EUROSCARF. The yeast strains were grown in Yeast Extract Peptone Dextrose (YPD) medium or synthetic complete (SC) medium with appropriate dropouts. All restriction enzymes, High-fidelity Phusion polymerase, Taq polymerase, T4 ligase and alkaline phosphatase (CIP) were obtained from New England Biolabs (Beverly, MA, USA). Oligonucleotides used in the present study are listed in Table S1. The *EGFP* gene was PCR amplified from plasmid pKT127 (EUROSCARF), digested with *BamHI/XhoI*, and inserted into plasmid pRS425Gal1, to yield pRS425Gal1-EGFP. Plasmid pRS425Cup1 was created by replacing the GAL1 promoter with the CUP1 promoter by *SacI-BamHI* digestion. Plasmid pRS425Cup1-EGFP was constructed in a similar way using pRS425Cup1 as the recipient vector. PAAS, CYP76AD1 and DOD genes were synthesized by GenScript or Genewiz. Plasmid pRS425-PAAS/Pha2 and pRS425-CYP76AD1/DOD were constructed by the golden-gate assembly method as previously described (20). All the plasmids and strains used in this study are listed in Table S2.

CRISPR/Cas9-mediated genome editing was carried out as previously described (21). The promoter regions of CTR1 and CUP1 were PCR amplified from the genomic DNA of BY4741. In brief, plasmid p415-GPD-Cas9 together with approximately 200 ng gRNA plasmid and 500 ng of homologous DNA donors was transformed into the yeast via electroporation at 1.6 kV. After recovery for approximately 2-3 h, cells were collected by centrifugation at 6000 *g* for 1 min, washed twice and resuspended in ddH_2_O. Appropriate amounts of cells were then spotted on SC agar plate with uracil and leucine dropped out. The plates were incubated at 30°C for three days, colonies were randomly picked and subjected to diagnostic PCR for verifying genome modifications. The detailed genotype information could be found in Table S2.

### Green fluorescence intensity measurement

For investigating the copper inducible expression of *EGFP* reporter gene, experiments were carried out in 14 ml tubes. Plasmid pRS425Gal1-EGFP was first transformed into strain JS-CR (BY4741 derivative with *P_CUP1_-Gal4* and *P_CTR1_-Gal80*). For the control, plasmid pRS425Cup1-EGFP was transformed into strain BY4741. Colonies were picked and inoculated into SC medium with leucine dropped out. Next day, 14 ml sterile tubes containing 2 ml SC-LEU medium supplemented with 2% glucose and different concentrations of copper sulfate (0, 0.5, 1.0, 2.5, 5.0, and 10 μM) were inoculated with fresh overnight cultures to an initial OD_600_ of 0.1. After 24 h cultivation, 100 μL of cell culture was taken for determining the cell density and green fluorescence intensities using the microplate reader Synergy H1 (BioTek, USA).

### Copper detection by betaxanthin-based colorimetric assay

For developing the betaxanthin colorimetric assay for the Cu(II) detection, plasmid pRS425-CYP76AD1/DOD was first transformed into strain JS-CR(2M), to give strain JS-BET. Colonies were picked from the plate and inoculated into SC medium with leucine dropped out. Next day, 14 ml sterile tubes containing 2 ml SC medium supplemented with 2% glucose and different concentrations of copper sulfate (0, 0.5, 1.0, 2.5, 5.0, and 10 μM) were inoculated with fresh overnight cultures to an initial OD_600_ of 0.1. oDA was added after 12 h cultivation to a final concentration of 0.5 mM when needed. The image of betaxanthin-producing yeast was captured after 24 h cultivation.

For monitoring the Cu(II) level in real water samples, the yeast cell suspension was mixed with real water samples and incubated in a 96-well plate at room temperature. 2 ppm Cu(II) was spiked into the real water samples to mimic the contaminated water. The color changes were captured after 24 h incubation.

### Development of 2-phenylethanol-based olfactory output

Plasmid pRS425-PAAS/Pha2 was first transformed into strain JS-CR(2M), to give strain JS-OLF1.0. 2 ml SC media (2% glucose + 2 mM L-phenylalanine) with different concentrations of copper sulfate (0, 0.5, 1.0, 2.5, 5.0, and 10 μM) was inoculated with fresh overnight cultures to an initial OD600 of 0.1. For the quantitation of 2-PE levels, the supernatant from cell broth was subjected to high-performance liquid chromatography (HPLC) analysis. Shimadzu Prominence LC-20A system (Shimadzu, Japan) was equipped with a photodiode array detector (DAD) under reversed-phase conditions. Mobile phase: 70% water with 0.1% trifluoroacetate + 30% acetonitrile. Column: Shimadzu C18 column (150 × 4.6 mm, 2.7 μm). Flow rate: 1 ml min^−1^. Column temperature: 40°C. Detection wavelength: 210 nm. The retention time for 2-PE is about 6.2 min.

### Development of styrene-based olfactory output

Plasmid pRS425-PAL2/FDC1 was first transformed into strain JS-CR(2M), to give strain JS-OLF2.0. Fresh overnight cultures were inoculated into 2 ml SC media (2% glucose + 2 mM L-phenylalanine) with different concentrations of copper sulfate (0, 0.5, 1.0, 2.5, 5.0, and 10 μM) to an initial OD600 of 0.1. 50% of dodecane was added after 12 h for the extraction of styrene from the fermentation broth. For the quantitation of styrene levels by gas chromatography – flame ionization detector (GC-FID), the organic phase was acquired by phase separation via centrifugation at 13,000 rpm for 10 min. The styrene peak was determined using Shimazu GC-2030 equipped with a Rtx-5 column (30 m × 250 μm × 0.25 μm thickness). Nitrogen (ultra-purity) was used as the carrier gas at a flow rate of 1.0 ml/min. GC oven temperature was initially held at 40°C for 2 min, increased to 45°C with a gradient of 5°C/min and held for 4 min. And then it was increased to 230°C with a gradient of 15°C/min. Authentic standard from Sigma-Aldrich was used for determining the styrene levels.

## RESULTS

### The design of yeast biosensor with reduced noise for Cu(II) detection

In *S. cerevisiae*, copper ion uptake is mediated by CTR1 and CTR3 encoded membrane-associated copper high-affinity transporters. Once inside the cell, the excess amount of copper ion is sequestrated in vacuoles (22) or by metal-binding proteins, such as metallothioneins of CUP1 and CRS5 (23). In *S. cerevisiae*, CUP1 is transcriptionally activated by Cu(II) via the copper-binding transcription factor Ace1 (24), whereas CTR1 and CTR3 are subjected to the copper-regulated transcriptional suppression mediated by the nutritional copper sensor of Mac1 (25, 26). Recently, copper-inducible expression based on the CUP1 promoter has been used for copper-induced expression of targeted proteins (27). However, the CUP1 promoter had a relatively high basal level because of the background copper level in the culture medium and only 20-fold of induction could be achieved upon the addition of Cu(II) (18, 19).

The galactose-inducible (GAL) system is one of the most tightly-regulated expression systems in yeast, which is subjected to glucose repression in the glucose-containing medium and de-repressed when galactose is used as the alternative carbon source (19, 28). GAL1, GAL7 and GAL10 mRNAs are rapidly induced >1000-fold on the addition of galactose (29). In this study, we attempted to reprogram the GAL system into a reduced-noise and ultrasensitive copper-sensing device. As depicted in Fig. 1A, a secondary genetic layer was introduced to control the key components involved in the GAL system, namely, the Gal4 activator and Gal80 repressor (30). To make GAL promoters respond to copper ion, the endogenous copper-repressible promoter CTR1 from yeast was used to control the Gal80 repressor, and the Gal4 activator was put under the control of the copper-inducible promoter CUP1.

**Figure 1.**
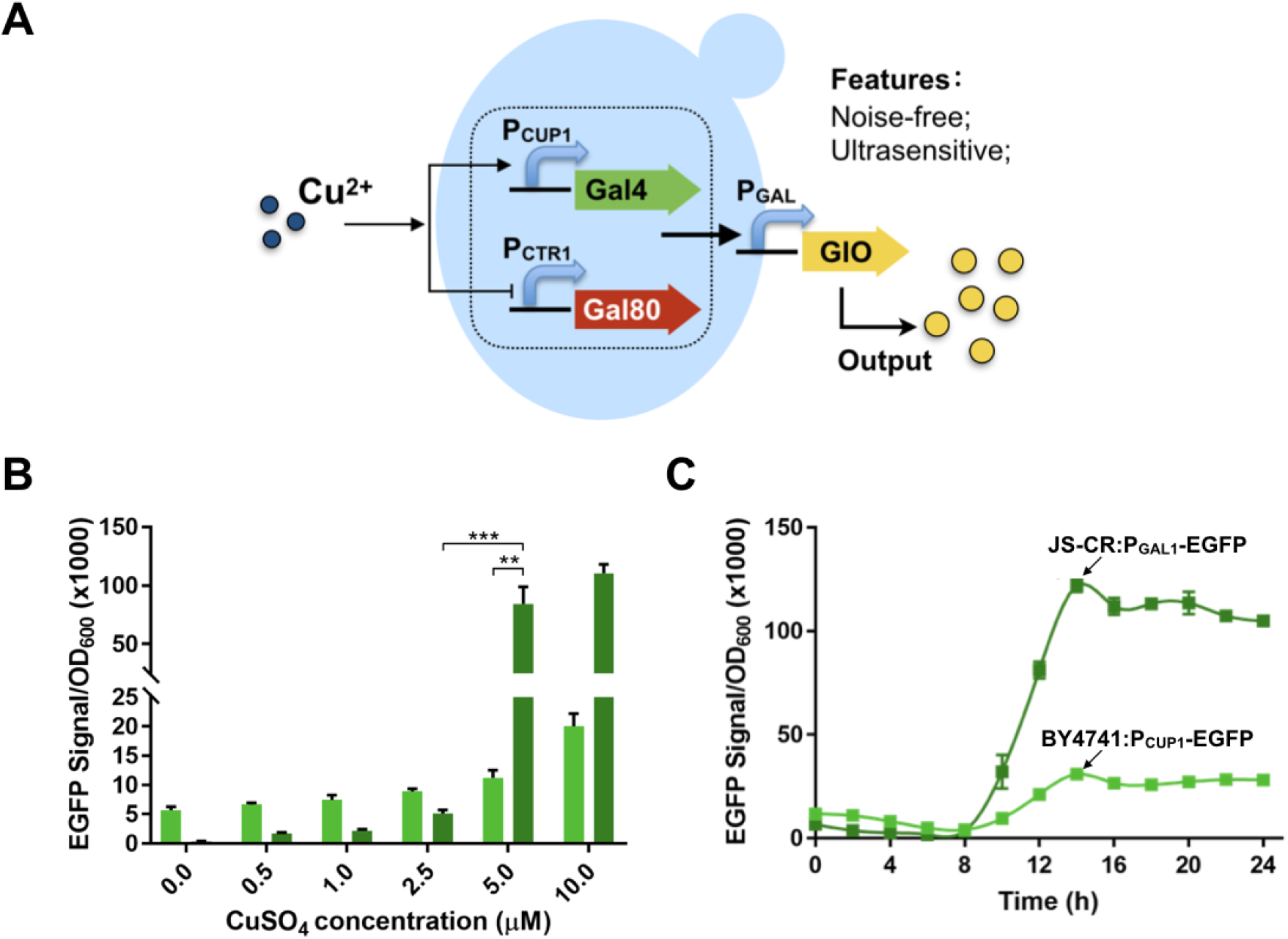
Frabrication of a reduced-noise and ultrasenstive copper-sensing circuit. (**A**) Reprogramming the galactose-inducible system into a copper-sensing circuit via a layered genetic design. Gal4, the activator in the galactose-inducible system; Gal80, the repressor from the galactose-inducible system; P_CUP1_, copper-inducible promoter; P_CTR1_, copper-repressible promoter; P_GAL_, galactose-inducible promoters. (**B**) EGFP signal output of BY4741 harboring pRS425Cup1-EGFP (light green bars) and JS-CR harboring pRS25Gal1-EGFP (dark green bars) in response to different concentrations of copper sulfate. Green fluorescence intensities were measured with excitation/emission at 476/512 nm. The data are collected after 24 h and represent the mean value with standard deviation. (C) Time course of EGFP levels in response to 10 μM copper sulphate. Experiments were carried out in triplicate. Statistical analysis was performed by use *t*-test (One-tailed, two-sample unequal variance; \*\**p* < 0.01, \*\*\**p* < 0.001).

### Characterization of the performance of yeast biosensor for Cu(II) detection

Although the Gal4 activator would be expressed at a certain amount due to the leakiness of P_CUP1_, the Gal80 repressor could keep the GAL system at its “OFF” state since the copper sensor of Mac1 triggers the expression of Gal80 repressor under the CTR1 promoter at the nutritional copper level. As shown in Fig. 1B, we successfully addressed the leaky problem for copper detection commonly encountered by other studies. BY4741 derived strain JS-CR harboring plasmid pRS425Gal1-EGFP (enhanced green fluorescent protein) nearly gave no appreciable EGFP signal at the nutritional copper level. When treated with 5 μM Cu(II) (~0.32 ppm), the genetic switch was turned on with a sharp increase of EGFP signal and >300-fold signal output was observed when 10 μM Cu(II) (~0.64 ppm) was added. In contrast, strain BY4741 with pRS425Cup1-EGFP (the conventional design) had a relatively high basal level of EGFP signal and only 4-fold signal output was achieved when exposed to 10 μM Cu(II) (Fig. 1B). As shown in Fig. 1C, the time course study of EGFP output in response to 10 μM Cu(II) revealed that the maximum output of our engineered yeast sensor could be reached around 14 h incubation.

These findings confirmed that the layered genetic circuit could effectively address the leaky issue encountered by the traditional designs, and the enhanced dynamic range of our biosensor could guarantee more reliable results with less chance of false positives. Besides, we also applied the yeast biosensor for detecting other heavy metals (Fig. S1). Interestingly, we found that no appreciable amount of EGFP signal was observed upon the addition of 10 μM Fe(III), Mn(II), Zn(II), Co(II), Cd(II), Ag(I) or Hg(II). Previously, Cd(II) and Hg(II) were documented to trigger the repression of CTR1 and CTR3 gene (31). However, Cd(II) and Hg(II) require concentrations 3 orders magnitude greater to down-regulate the CTR3 expression. We reasoned that the synthetic circuit is at the “OFF” state due to insufficient repression of Gal80 expression upon the exposure of 10 μM Cd(II) and Hg(II), thereby making the yeast sensor relatively specific toward the Cu(II) detection.

### Copper ion detection via the betaxanthin-based colorimetric assay

We next sought to create a transformative on-site copper detection device unattainable by traditional methods. As shown in Fig. 2A, we devised a betaxanthin-based colorimetric assay for potential field-deployable copper detection. Heterologous expression of CYP76AD1 from sugar beet *Beta vulgaris* and L-DOPA dioxygenase (DOD) from *Mirabilis jalapa* has been used for the betaxanthin production in budding yeast (32, 33). As can be seen from Fig. 2B, strain JS-BET with CYP76AD1 from *B. vulgaris* and DOD from *M. jalapa* gave noticeable orange colors when the cells were exposed to >5 μM Cu(II) (~0.32 ppm). The visual effect could be substantially enhanced by adding 0.5 mM amino donors such as *o*-dianisidine (oDA) (Fig. 2B). Therefore, our engineered yeast sensor confers a narrow range switch-like behavior, which can give a “yes/no” response from the color changes.

**Figure 2.**
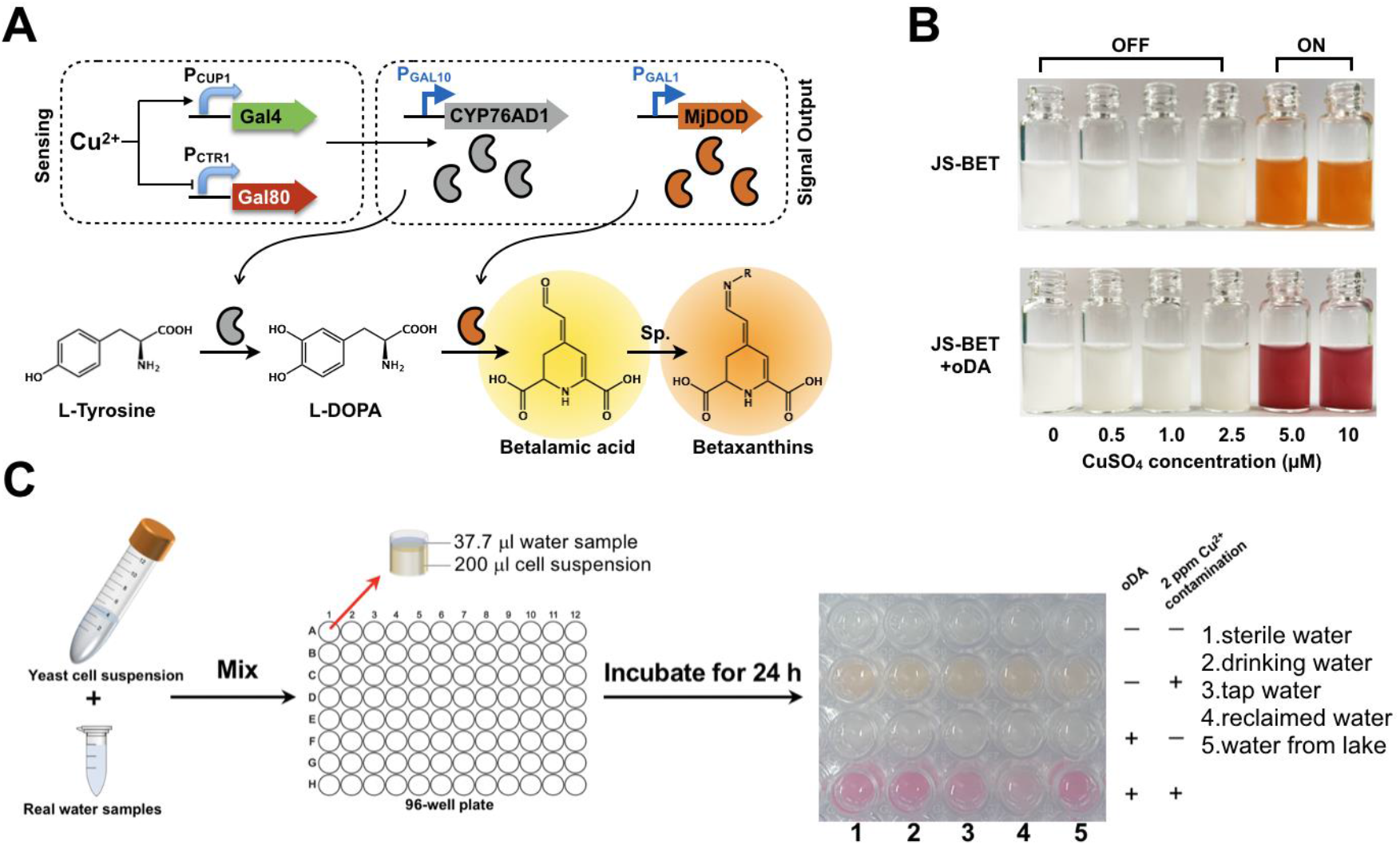
Copper ion detection by the betaxanthin-based visual phenotype. (**A**) Schematic description of copper detection by the product of betaxanthin in budding yeast. L-DOPA, L-3,4-dihydroxyphenylalanine; CYP76AD1, cytochrome P450 tyrosine hydroxylase from *B. vulgaris*; MjDOD, L-DOPA dioxygenase from *M. jalapa*. (**B**) Copper detection by betaxanthin-based colorimetric assay using strain JS-BET. Different concentrations of copper sulfate (0, 0.5, 1.0, 2.5, 5.0, and 10 μM) were added to the culture medium to mimic the copper-contaminated water. The image of betaxanthin-producing yeast was captured after 24 h cultivation. (**C**) Potential on-site detection of copper ion in different water samples. Different sources of water samples with or without 2 ppm copper ion contamination were tested.

According to the world health organization (WHO), the Cu(II) level above 2 ppm is considered to be hazardous to humans and the environment (34). As a proof-of-concept, we further attempted to develop a 96-well plate based colorimetric assay for monitoring the Cu(II) contamination in real water samples. In brief, the freeze-dried yeast cells were resuspended in the cell broth. Approximately 1:6 (volume) of water to yeast suspension were mixed together and incubated at the room temperature for 24 h. As shown in Fig. 2C, the control samples did not produce noticeable colors, whereas all contaminated water samples with 2 ppm Cu(II) gave the expected color changes. Noteworthily, we found that the reclaimed water sample showed a less intense color, indicating that there might be some other chemicals affecting the performance of our yeast sensor. Therefore, future work will be required to identify these interfering factors before the yeast biosensor can be used for practical applications.

### Copper ion detection based on olfactory outputs

*S. cerevisiae* that capitalizes on the orthogonality and specificity of its G protein-coupled receptor (GPCR) mating pathway has been engineered toward an “olfactory yeast” and the resulting yeast could detect an explosive residue mimic of the odorant 2,4-dinitrotoluene (35). More recently, an olfactory yeast biosensor that detects the hormone estradiol signal based on an odor product of isoamyl acetate was reported (36). As inspired by these “olfactory” yeasts, we also attempted to develop odor-based olfactory outputs for Cu(II) detection. 2-Phenylethanol (2-PE) is a fragrant compound that gives a rose-like smell. *S. cerevisiae* naturally synthesizes 2-PE as the fusel alcohol via the Ehrlich pathway (37). To reduce the background 2-PE level from the endogenous Ehrlich pathway, *S. cerevisiae* with Δ*aro10* to attenuate the Ehrlich pathway (38) was used as the starting strain. Next, a heterologous phenylacetaldehyde synthase (PAAS) from *Petunia* (39) was used for 2-PE production (Fig. 3). As shown in Fig. 3A, the background 2-PE level from the Ehrlich pathway could reach 20.93 ± 1.79 mg/L, whereas 119.69 ± 11.05 mg/L of 2-PE was obtained upon the addition of 10 μM Cu(II) for the yeast biosensor equipped with the copper-sensing genetic circuit. However, we could not clearly distinguish the samples contaminated by copper based on a blind test between the lab members.

**Figure 3.**
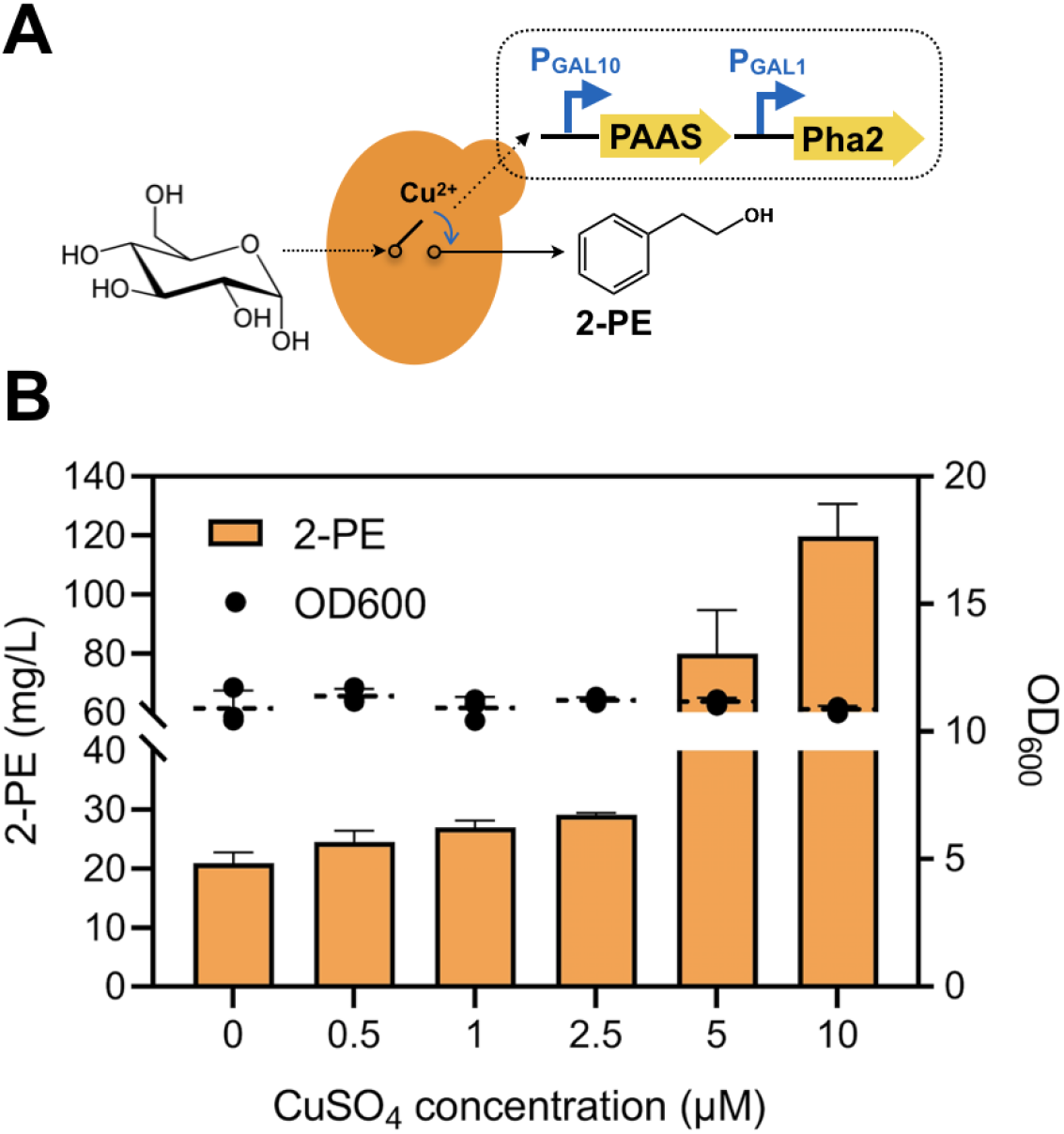
Copper ion detection based on the olfactory output of 2-phenylethanol. (**A**) Biosynthetic route toward 2-PE synthesis. PAAS, phenylacetaldehyde synthase from *Petunia;* Pha2, prephenate dehydratase from *S. cerevisiae*. (**B**) The 2-PE levels in response to different concentrations of copper sulfate (0, 0.5, 1.0, 2.5, 5.0, and 10 μM). The 2-PE concentrations were measured after 24 cultivation. All experiments were performed in triplicate and the data represent the mean value with standard deviation.

To further create a reduced-noise olfactory output for Cu(II) detection, we next used the odor product of styrene that is not naturally produced by *S. cerevisiae* for a proof-of-concept study (Fig. 4). Phenylalanine ammonia-lyase (PAL2) from *Arabidopsis thaliana* and ferulic acid decarboxylase (FDC1) from *S. cerevisiae* were used to convert L-phenylalanine into styrene (38). As shown in Fig. 3B, the second-generation design of styrene-based olfactory output solved the problem of the background odor, and 21.0 ± 1.48 mg/L of styrene was obtained upon the addition of 10 μM Cu(II), which is above the threshold of human olfactory system.

**Figure 4.**
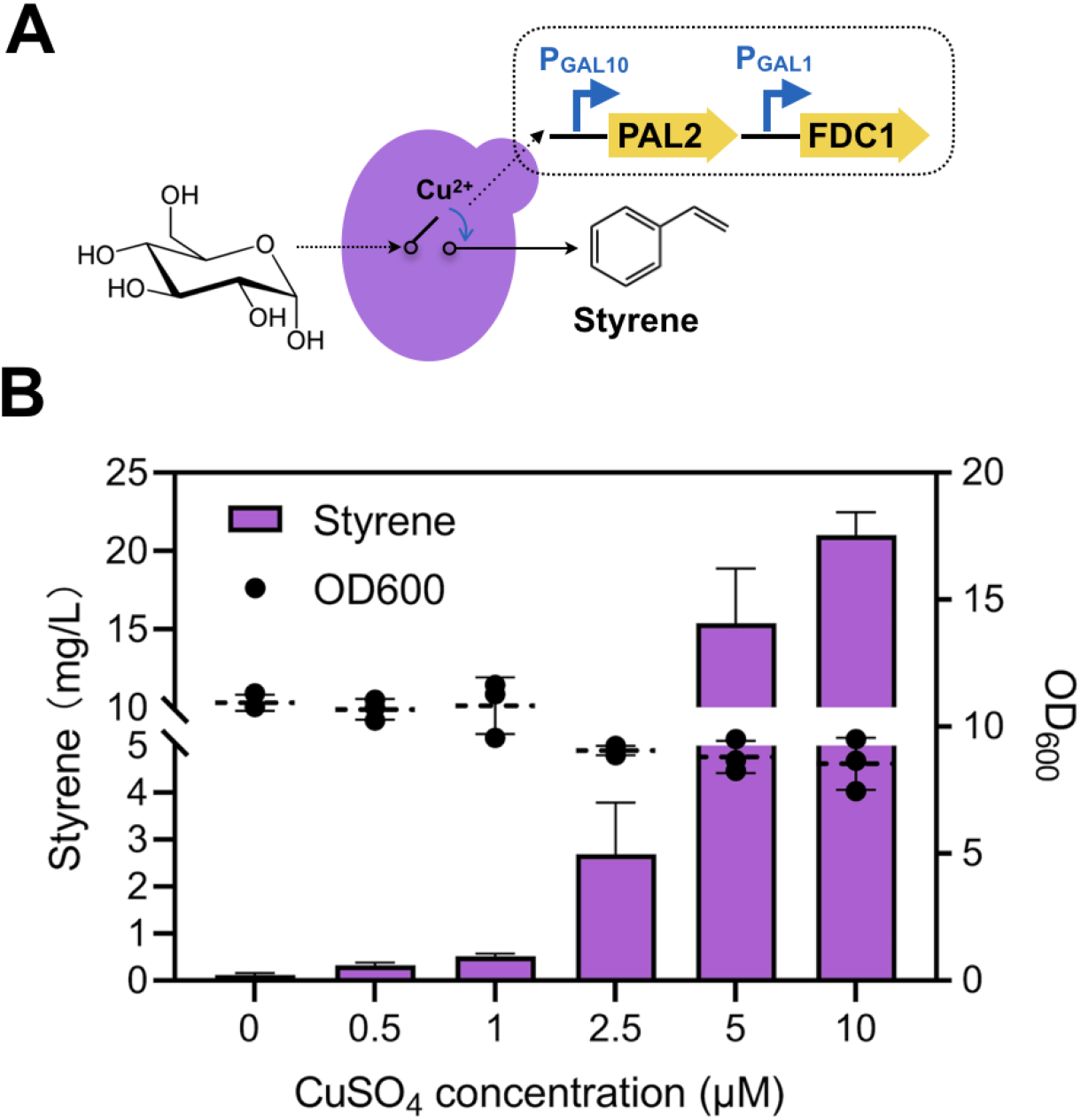
Copper ion detection based on the olfactory output of styrene. (**A**) Biosynthetic route toward styrene synthesis. PAL2, phenylalanine ammonia lyase from *A. thaliana*; FDC1, ferulic acid decarboxylase from *S. cerevisiae*. (**B**) The styrene levels in response to different concentrations of copper sulfate (0, 0.5, 1.0, 2.5, 5.0, and 10 μM). The styrene concentrations were measured after 24 cultivation. All experiments were performed in triplicate and the data represent the mean value with standard deviation.

## DISCUSSION

Nowadays, biosensors are no longer just a combination of microorganisms and physical transducer for detecting specific signals (40). Although a number of yeast biosensors have been developed for the detection of heavy metals and other environmental pollutants (14), many limitations remain with regard to their performance for on-site testing and accuracy. In this study, we have demonstrated for the first time to reprogram *S. cerevisiae* into a low-noise and reliable biosensor that permits potential field-deployable Cu(II) detection. By refactoring the tightly-regulated GAL system into a copper-sensing device, we not only addressed the issue of leaky expression caused by the nutritional copper ion level but also enhanced the magnitude of signal output to >300-fold.

At this moment, ~0.32 ppm Cu(II) can give a clear signal output by our yeast-based biosensor, which is similar to the microfluidic paper-based technique (9). For potential on-site copper detection, we have developed the betaxanthin-based colorimetric assay which could give a “yes/no” response from the color changes. Furthermore, we also investigated olfactory outputs based on 2-phenylethanol and styrene for the Cu(II) detection. Future efforts will be required to further improve the response time of the synthetic circuit as well as increase its sensitivity, before the yeast sensors can be used for real-world applications.

## Author Contributions

J.Y. conceived of the project and wrote the paper. J.Y. and C.F. constructed all the plasmids and strains. C.F., D.Z., and Q.M. collected the data. The authors would like to thank the lab members for blind-testing all the experiments.

## Funding

This work was supported by Xiamen University (grant no.: 0660-X2123310) and ZhenSheng Biotech, China.

## Compliance with ethical standards

### Conflicts of interest

The authors declare that they have no competing interests.

### Ethical approval

This study does not contain any studies with human participants or animals performed by any of the authors.

### Data availability

All data generated or analysed during this study are included in this published article [and its supplementary information files].

## TOC

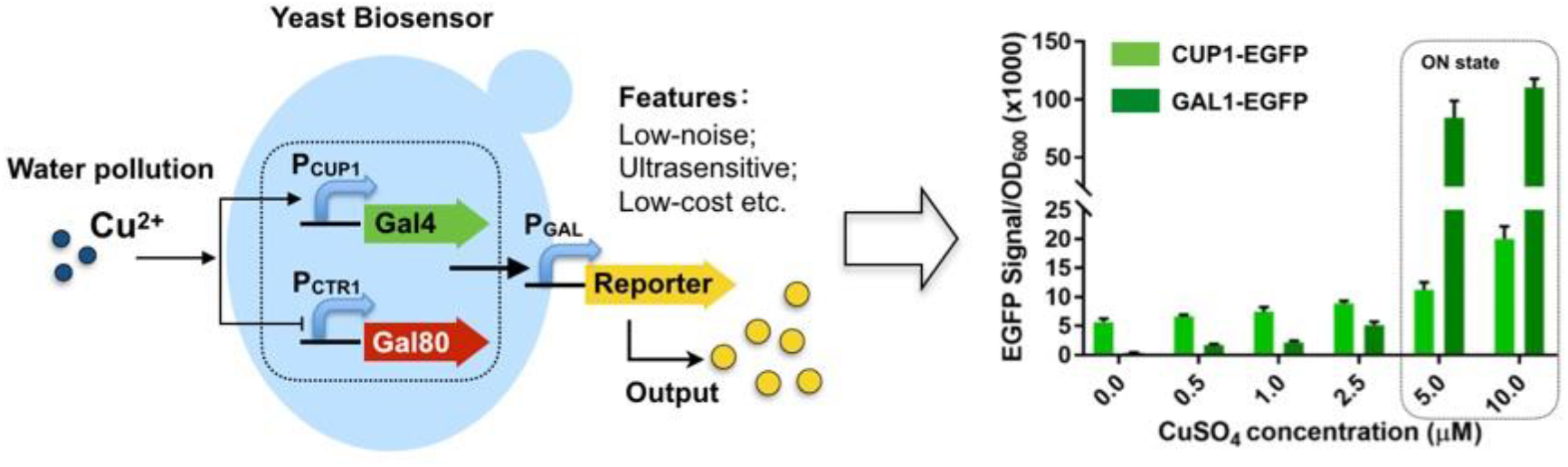

